# Distinct causes underlie double-peaked trilobite morphological disparity

**DOI:** 10.1101/2024.01.05.574297

**Authors:** Harriet B. Drage, Stephen Pates

## Abstract

Trilobite cephalic morphology impacted the autecology of individuals, and is critical for high- and low-level taxonomic assignments. Disparity in trilobite cephalon shape varied through time and was integral to the occupation of a diversity of ecological niches. To fully appreciate trilobite cephalic evolution, we must understand how this disparity varies, and what factors control cephalon morphometry. We explore the disparity of trilobite cephala through the Palaeozoic, and analyse the associations between cephalic morphometry and taxonomic assignment and geological Period, using a dataset of 983 2D trilobite cephalon outlines. Elliptical Fourier transformation visualised as a Principal Components Analysis suggests significant differences in morphospace occupation for order and Period groups, and comparisons of disparity measures also suggest significantly different disparities between the groups. Trilobite cephalic disparity peaks in the Ordovician and Devonian. The Cambrian–Ordovician expansion of morphospace occupation appears a result of radiation to new niches, and thus all trilobite orders were established by the late Ordovician. In comparison, the morphospace expansion from the Silurian to Devonian seems solely a result of within-niche diversification rather than novel niche occupation. However, analyses interrogating the regions of morphospace occupied, including centroid distances, average pairwise shape comparisons and Linear Discriminant Analysis demonstrate that, except for the order Harpida and the Cambrian and Ordovician Periods, order and geological Period could not be robustly predicted for an unknown trilobite. Further, Kmeans clustering analyses suggest the total dataset naturally subdivides into only seven groups that do not correspond with taxonomic orders, though Kmeans clusters do decrease in number through the Palaeozoic, aligning with findings of decreasing disparity.

## Introduction

The fossil record documents the origins and radiations of animal groups and their morphological variation. Critically, the fossil record provides a perspective of how life on Earth has changed through time, including abundant information on long-extinct groups, allowing inferences about the impacts of environmental change, extinction events, and radiations on the diversity and morphological disparity of animal groups. This morphological disparity, which is often decoupled from taxonomic diversity (e.g., Bapst et al., 2012; Fortey et al., 1996; Puttick et al., 2020; Wan et al., 2021; Foote 1993a; Hopkins 2013), results from interlinked ecological, functional and taxonomic components (e.g., Foote 1997; Cantalapiedra et al., 2017; Hopkins, 2014). Disparity is often quantified through morphometry and visualised as morphospaces, with relative trends in occupation area and location through time for clades informing on the patterns and processes that impacted their morphological variation on geological timescales. Increases or decreases in the volume of morphospace occupied (often quantified through metrics such as sums of ranges, and sums of variances; e.g., Lloyd, 2016; Guillerme et al., 2020a) indicate a corresponding increase or decrease in the morphological disparity of a group, often caused by extinction or radiation events lacking selectivity or acting only at the margins of the morphospace (e.g., Korn et al., 2013). A complementary metric, the change in position of the centroid in morphospace through time (or average position in the morphospace), indicates a change in the mean shape of members of the group, which can be interpreted as selective extinctions and radiations favouring particular morphologies, likely linked to an ecological mode or environmental factor (Korn et al., 2013).

Trilobites are one of the most abundant animal groups preserved in the fossil record, owing to their notable global diversity across the Palaeozoic, their strongly biomineralised exoskeletons with high preservation potentials, and their colonisation of a range of marine environments worldwide (e.g., Adrain et al., 1998; Brandt, 2002; Esteve et al., 2021; Fortey, 2014, 2001; Whittington et al., 1997; Fig. 1). Thus, they provide an excellent study group with which to explore the interrelated temporal (including changing ecological and functional pressures) and taxonomic components acting on the morphological disparity of a diverse clade. Trilobites originated during the early Cambrian, with the oldest body fossils of this group c. 521 million years old (e.g., Paterson, 2019). Analyses of their raw diversity have revealed two major peaks in the global record, the largest in the middle Cambrian (Webster, 2007) and the second in the early Devonian (Adrain, 2008; Bault et al., 2022), before a dramatic reduction during the late Devonian (e.g., Bault et al., 2023a; Lerosey-Aubril and Feist, 2012), leaving only one major extant order (Proetida, plus Aulacopleurida if recognised as an order) with reduced diversity by the Carboniferous and a final extinction event at the end Permian 252 Ma (e.g., Lerosey-Aubril and Feist, 2012). While the middle Cambrian peak in diversity correlates with a peak in intraspecific variation for the group (Webster, 2007), previous studies have differed on when the peak in trilobite disparity occurred. Earlier studies assessing changing disparity over geological time scales focused on the outline shape of the cranidium, and found a disparity peak in the Ordovician (Foote, 1993b, 1991), facilitated by environmental preferences of taxa (Hopkins, 2014). However, a more recent study using the outline of the whole cephalon (cranidium and librigenae), albeit with a significantly smaller sample size (400 compared to 1125 in Foote, 1993b), found an earlier Cambrian peak in disparity and a second peak in the Devonian, broadly mirroring reported diversity trends (Suárez and Esteve, 2021).

**Figure 1:**
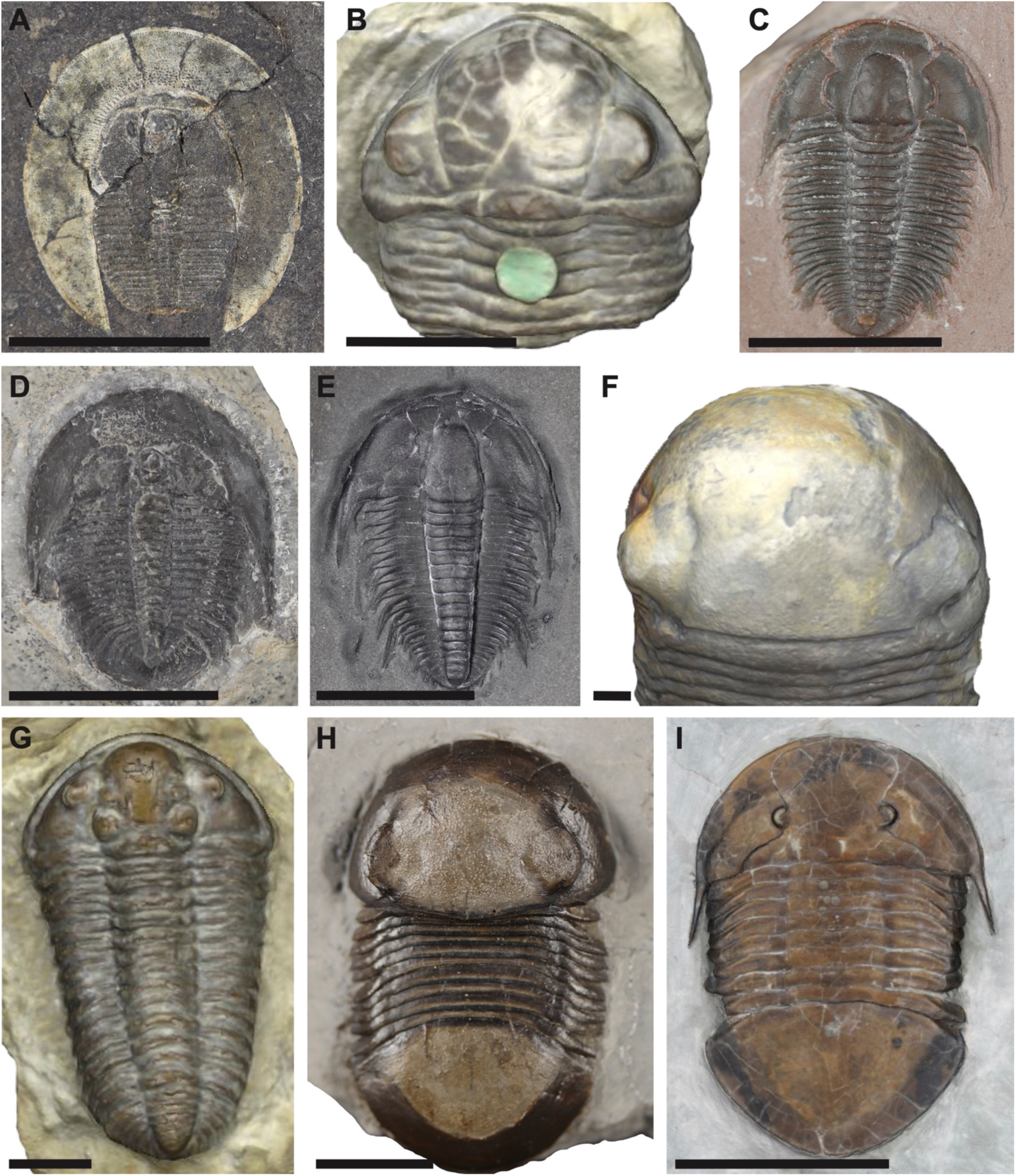
Examples of trilobite specimens included within the dataset. A, *Bohemoharpes naumanni* Senckenberg Museum X286a; B, snapshot of 3D model, *Acaste downingiae* NHMUK 44409; C, *Modocia typicalis* Senckenberg Museum unnumbered Westphal collection; D, *Pterocephalia norfordi* Senckenberg Museum unnumbered Westphal collection; E, *Orygmaspis contracta* Senckenberg Museum unnumbered Westphal collection; F, snapshot of 3D model, *Bumastus barriensis* NHMUK I1029; G, snapshot of 3D model, *Calymene blumenbachii* NHMUK 44215; H, *Bumastus iorus* Senckenberg Museum unnumbered Westphal collection; I, *Megalaspides* sp. SMNKPAL 3887. Scale bars = 1 cm.

While morphometric analyses on trilobite data have been used to address a range of macroevolutionary (e.g., Foote 1989, 1991, 1993a, b; Hopkins 2014; Suárez & Esteve 2021), behavioural (e.g., Drage, 2022; Drage et al., 2023; Suárez and Esteve, 2021), developmental (e.g., Crônier et al., 1998; Hopkins and Pearson, 2016; Kim et al., 2002), systematic (e.g., Holmes et al., 2020; Martin et al., 2023; Paterson, 2005; Żylińska et al., 2013), and taphonomic (e.g., Hopkins and Pearson, 2016; Webster and Hughes, 1999) questions, several crucial macroevolutionary questions remain.

1. What are the extents of trilobite disparity in cephalic morphometry?
2. How does cephalic morphometry vary with order-level taxonomy, and can cephalon shape be used to predict taxonomic assignment?
3. How does cephalic morphometry vary across the Palaeozoic, in terms of morphospace volume occupation and movement through this space, and can cephalon shape be used to predict the geological Period occupied?

To address these questions, we analyse a dataset of nearly 1000 trilobite cephala, digitised as 2D outlines. We quantify the taxonomic (ordinal level) and temporal changes in morphospace occupation for Trilobita, documenting how the morphology of this key Palaeozoic group changed over the course of 250 million years. We also provide all data and R code (https://osf.io/vz9a5/), and the analytical protocol for these analyses.

### Previous works on the disparity of Trilobita or groups therein

There is an extremely rich history of both traditional and geometric morphometric approaches aimed at furthering our understanding of varied aspects of trilobite evolution, across taxonomic scales. In fact, the wealth of these studies as applied to specific trilobite species or groups are too numerous to fully cover outside of a dedicated review (e.g., Abe and Lieberman, 2012; Bordonaro et al., 2013; Crônier et al., 2005, 2004, 1998; Crônier and Fortey, 2006; Delabroye and Cronier, 2008; Foote, 1997, 1993, 1991; Gendry et al., 2013; Gerber and Hopkins, 2011; Hong et al., 2014; Hughes and Chapman, 1996, 1995; Labandeira and Hughes, 1994; Park et al., 2008; Rábano et al., 2008; Sheets et al., 2004; Smith and Lieberman, 1999; Webster, 2020, 2011; Webster and Zelditch, 2011; Zhao et al., 2020). The use of elliptical Fourier analyses to quantify the disparity of trilobite cranidia was pioneered by Foote (1989). This dataset was then expanded (e.g., Foote 1991, 1993a, b) and used to explore links between environmental and morphological characteristics within the group (Hopkins 2014). More recently, studies have used morphometric data pertaining to the whole cephalon. A number of key recent studies following these are discussed, though as these examples and the references therein demonstrate, a full review of trilobite morphometric analyses is beyond the scope possible here.

Several recent studies have quantified trilobite cephalic morphometry using various protocols and large-scale datasets to explore their diversity and disparity through geological time, and their potential links to other characteristics. Suárez and Esteve (2021) used 400 cephalon outlines (including agnostids) to assess trends in cephalic morphometry with the prevalence of different enrolment strategies through the Palaeozoic. Bault et al. (2023a) used a large dataset of cephala and pygidia across trilobite groups to determine whether disparity trends followed diversity across the Devonian. Holmes (2023) assessed cephalic disparity during the Cambrian, with different cephalic structures suggesting differing patterns in morphospace expansion. Cephalic outline showed rapid diversification in form, while interior structures showed a more gradual increase in disparity (Holmes, 2023). Hopkins et al. (2023) analysed diversity, disparity, and geological range in Permian trilobites and compared this to the other Palaeozoic Periods.

Many other studies have used morphometric analyses to explore the diversity and disparity of specific trilobite groups. These analyses have been used to assess shape change during ontogeny, such as for *Cryptolithus tesselatus* Green, 1832 (Hopkins and Pearson, 2016), to better understand development trajectories. Several studies explored shape evolution within specific clades; for example, Hopkins (2017) coupled cranidial shape change with exoskeleton morphological characters in pterocephaliids, and Bault et al. (2023b) explored shape evolution in Phacopidae and the associations between disparity and external forcing events. Many studies have used morphometric analyses to aid in taxonomic work (e.g., Álvaro et al. 2018; Zhao et al. 2020). These analyses can also be informative for building or testing phylogenetic hypotheses; for example, Martin et al. (2023) produced a phylomorphospace for asteropyginid glabellae. Still more research used trilobite morphometric data to explore specific morphological or behavioural evolutionary questions. For example, Vargas-Parra and Hopkins (2022) tested patterns of modularity in the trilobite cephalon, and hypotheses relating to the developmental origins of the eye. Drage (2022) used traditional multivariate morphometric analyses to test for an association between body proportions and exoskeleton moulting behaviour across Trilobita, while Drage et al. (2023) tested the same hypothesis on an intraspecific scale, using a large dataset of *Estaingia bilobata* Pocock, 1964.

## Methods and materials

### Using the cephalon as a proxy for trilobite disparity

A large part of trilobite diversity and disparity of forms is manifested in their cephalic morphometries; the shapes and sizes of the cephala of different groups. The variation in functional morphology of the cephalon is intrinsically linked to the varied life modes of trilobites; different cephalic shapes and structures have been suggested to be adaptations to different feeding modes (e.g., Fatka and Szabad, 2011; Fortey and Gutiérrez-Marco, 2022; Fortey and Owens, 1999; Hegna, 2010; Hughes, 2000; Pearson, 2017), life modes (e.g., Bault et al., 2023b; Cherns et al., 2006; Esteve et al., 2021; Fortey, 2014), or specific behaviours (e.g., Drage, 2019; Henningsmoen, 1975; Suárez and Esteve, 2021). However, the evolution and extent of disparity of the trilobite cephalon remains unclear, with uncertainty around the unstable high-level trilobite taxonomy (Adrain, 2013, 2011; Paterson, 2019), the potential homology of cephalic structures (Du et al., 2023; Hughes, 2003; Park and Kihm, 2017), and the adaptation of cephalic shape to hypothetical life mode.

While cephalic outline does not represent the whole (Holmes, 2023), the lack of homologous morphological points otherwise shared between all trilobite groups makes this a way to utilise the broadest taxonomic dataset. Furthermore, while 3D data might be expected to be more representative of form, previous studies comparing the use of 2D and 3D morphometric data on comparable scales suggest 2D data is sufficient for evolutionary hypothesis testing, as it reconstructs the same patterns as 3D data (Hopkins and Pearson, 2016). Compaction and flattening of specimens preserved in shales does increase the variance of landmark positions, alongside abaxial and anterior splaying of genal regions (Webster and Hughes, 1999), however, not on the same scale as interspecific differences in cephalic shape.

### Dataset construction

Photographs of 597 trilobite cephala, plus 386 additional specimens published by Suárez & Esteve (2021; with agnostids removed), representing at least 520 species total were gathered from the descriptive literature, online museum databases accessed through iDigBio, and additional museum photography (for references, search terms, and dates of access, see Appendices I and II). Specimens were photographed from the following museum collections: the Senckenberg Museum (183 specimens, representing 122 species), Sedgwick Museum, Cambridge (62 specimens, representing 46 species), Staatliches Museum für Naturkunde Karlsruhe (10 specimens, representing 9 species), and Natural History Museum, London (112 specimens, representing 81 species). Only reasonably intact cephala, without extreme deformation and with at least one librigena in place, were included; singular missing librigenae were reconstructed based on the shape and location of the present librigena. Work by Webster and Hughes (1999) suggested that flattening has a consistent effect on the morphometric data representing different species, and so that flattening deformation would likely have a minor impact on the results.

Cephalic outline semilandmark data for specimens were obtained using tpsDIG2 (Rohlf, 2015). For all specimens two curves were created: 1) from the tip of the left genal spine (or, in the absence of a genal spine, the posterior-left margin of the cephalon) clockwise around the anterior of the cephalon to the tip of the right genal spine (or, in the absence of a genal spine, the posterior-right margin of the cephalon); 2) from the tip of the right genal spine (or in the absence of a genal spine, the posterior-right margin of the cephalon) around the posterior of the cephalon to the tip of the left genal spine (or, in the absence of a genal spine, the posterior-left margin of the cephalon). Each curve consisted of 64 evenly spaced semilandmarks, giving a total cephalic outline for each specimen constructed of 128 semilandmarks (see Fig. 2). The data published by Suárez and Esteve (2021) were resampled to give an equivalent number of outline semilandmarks for all specimens.

**Figure 2:**
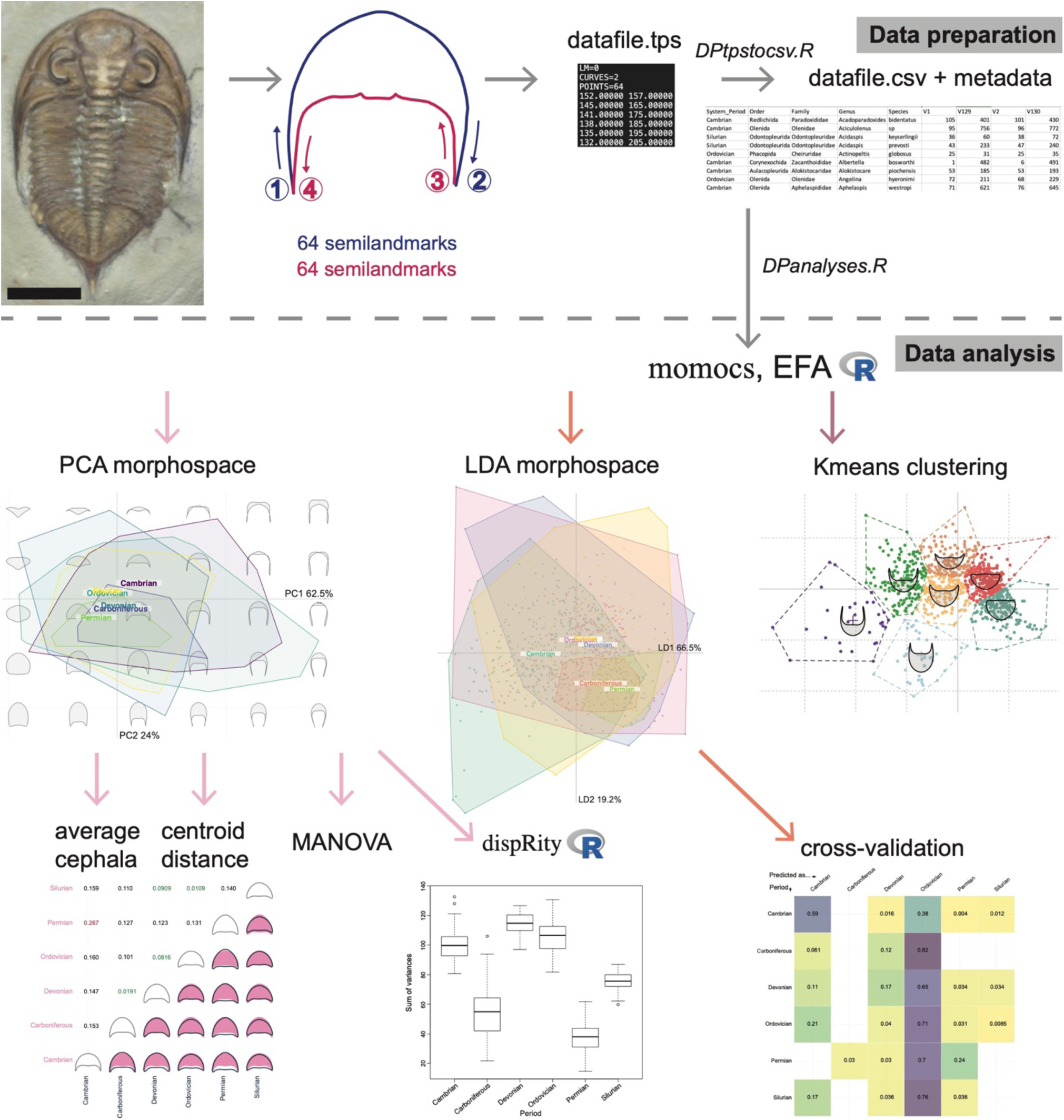
Cephalic outline collection (data preparation phase) and analysis (data analysis phase) methodology. Numbers and arrows on the cephalon outline show the process of producing two semilandmark curves to capture the cephalon shape. Filenames given reflect the R code to run the various steps provided at [https://osf.io/vz9a5/]. EFA, elliptical Fourier transformation; PCA, principal components analysis; LDA, linear discriminants analysis. Momocs R package by Bonhomme et al. (2014), dispRity R package by Guillerme (2018). Trilobite image is a snapshot of a 3D model made of *Dalmanites caudatus*, scale bar = 1 cm.

Metadata for all species included order assignment following Adrain (2011) and geological Period occupied following Jell and Adrain (2002). Groups traditionally included in the ‘Ptychopariida’ were grouped as ‘Unassigned’, due to the problematic nature of the clade (Adrain, 2011; Fortey, 2001; Suárez and Esteve, 2021). See Tables 1 and 2 for a summary of the dataset composition.

**Table 1:** PCA morphospace areas occupied by the convex hulls of the taxonomic order groupings, and calculated disparity measures (sums of variances and sums of ranges; plotted in Fig. 6).

| **Taxonomic order** | **Sample size (specimens)** | **Convex hull area (no unit)** | **Sums of variances (no unit)** | **Sums of ranges (no unit)** |
| --- | --- | --- | --- | --- |
| Asaphida | 115 | 0.31 | 0.065 | 1.534 |
| Aulacopleurida | 34 | 0.19 | 0.036 | 1.220 |
| Corynexochida | 68 | 0.29 | 0.045 | 1.425 |
| Harpida | 33 | 0.28 | 0.038 | 1.444 |
| Lichida | 21 | 0.16 | 0.048 | 1.089 |
| Odontopleurida | 23 | 0.18 | 0.043 | 1.029 |
| Olenida | 68 | 0.55 | 0.080 | 1.818 |
| Phacopida | 359 | 0.48 | 0.037 | 1.674 |
| Proetida | 100 | 0.25 | 0.034 | 1.211 |
| Redlichiida | 69 | 0.44 | 0.053 | 1.756 |
| Trinucleida | 38 | 0.51 | 0.133 | 2.023 |
| Unassigned | 55 | 0.31 | 0.043 | 1.436 |

**Table 2:** PCA morphospace areas occupied by the convex hulls of the geological Period groupings, and calculated disparity measures (sums of variances and sums of ranges; plotted in Fig. 9).

| **Geological Period** | **Sample size (specimens)** | **Convex hull area (no unit)** | **Sums of variances (no unit)** | **Sums of ranges (no unit)** |
| --- | --- | --- | --- | --- |
| Cambrian | 249 | 0.64 | 0.055 | 1.983 |
| Ordovician | 351 | 0.74 | 0.064 | 2.217 |
| Silurian | 169 | 0.40 | 0.047 | 1.556 |
| Devonian | 148 | 0.59 | 0.078 | 1.930 |
| Carboniferous | 33 | 0.14 | 0.029 | 0.995 |
| Permian | 33 | 0.096 | 0.021 | 0.831 |

### Analyses

Geometric morphometric analyses and multivariate statistical analyses were performed to test the following null hypotheses:

1. Trilobite taxonomic orders show no difference in their cephalic outline morphometries.
2. There were no broad-scale changes in cephalon morphometry through geological time. The data gathering and analytical protocol is presented in Figure 2 and described below.

Firstly, a total dataset morphospace was produced using elliptical Fourier analysis (EFA), with 13 harmonics that represent 99.9% of the variation retained. Principal Components Analyses (PCA) were then used to visualise relevant patterns of shape variation and analyse differences in morphospace occupation between groupings, including taxonomic groups and geological age groups. The PCA groupings were further interrogated by calculating the convex hull areas, and comparing the centroid distances between groups. MANOVA tests were performed to test for differences in the distributions of principal components (PC) scores across the different groupings, and further pairwise tests used to interrogate the PC score differences (alpha level of 0.05 with Bonferroni correction = 0.05/n pairings). Only PCs 1 and 2 were plotted and used for the subsequent analyses and interrogations presented here; the morphospace scree plots (Fig. 3) show that these are responsible for 62.5% and 24% of the variation respectively. PC3 represents 4.2% of the morphospace variation; on visualising the outline changes across PC3 it is clear this results from sample noise likely due to minor specimen compression and distortion, and so PCs from 3 onwards were discarded.

**Figure 3:**
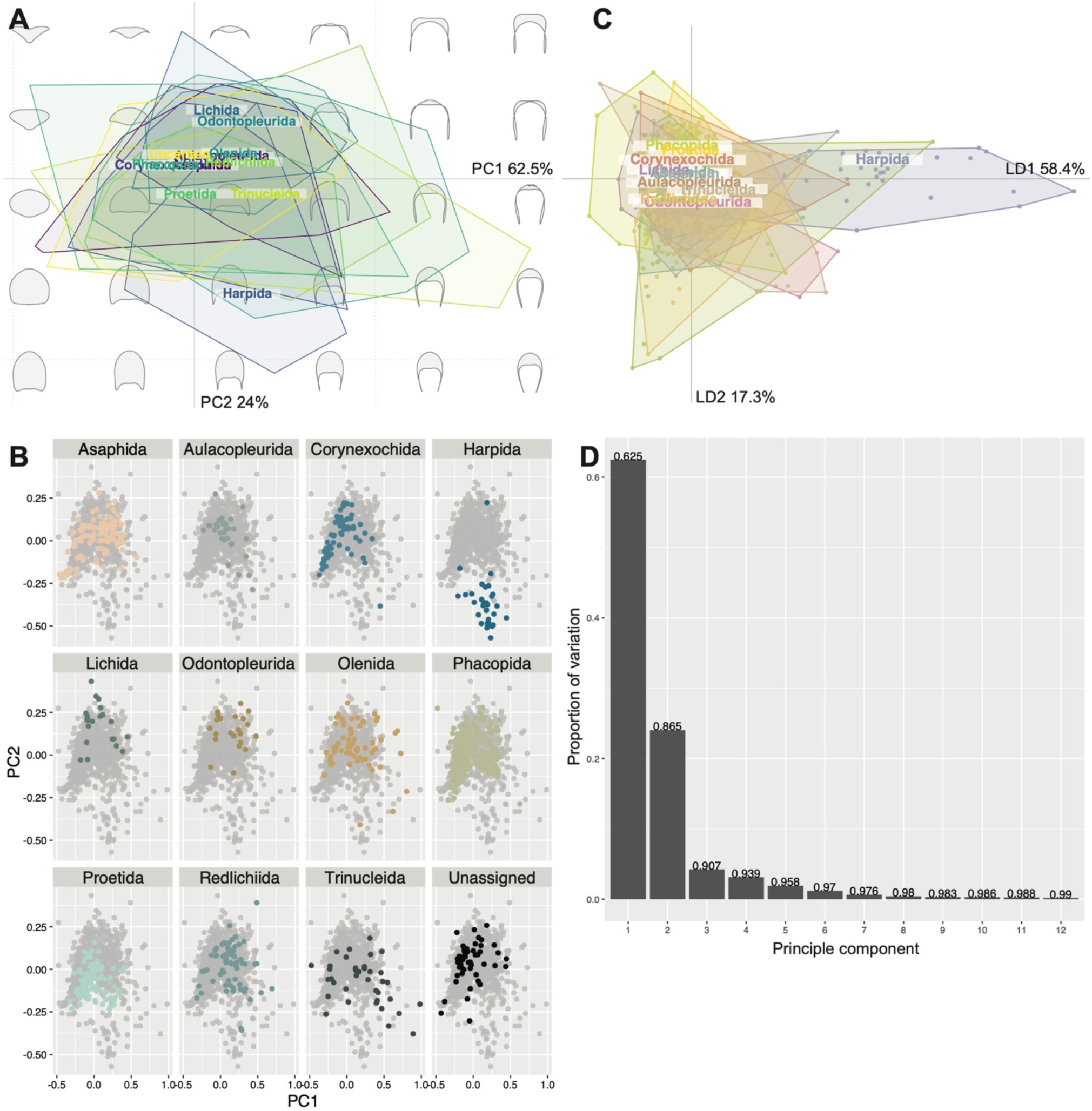
Principal components analysis (PCA) total plot (A), PCA facet wrap plot (B), and linear discriminants analysis (LDA) plot (C) of trilobite cephalic outline morphometry, grouped by taxonomic order. Principal components scree plot (D) is applicable to the entire dataset, and reflects no grouping. A and B visualise the same data, but B has the overlapping convex hull polygons separated out onto individual morphospaces.

Linear Discriminant Analyses (LDA) were also performed to visualise morphospace occupation when differences were maximised between the groupings. LDA cross-validation tables were produced to provide an alternative method of exploring the LDA overlap of groupings. These cross-validation tables inform on the probability that a new data point within each group would be incidentally placed within each other group, and therefore give an estimation of the predictiveness of cephalic outline for each grouping in the dataset.

Kmeans clustering analyses were performed to explore how cephalon morphometry stochastically grouped together in EFA morphospace. Mean cephalon outline shapes were produced for the groupings to visualise any differences in cephalic morphometry across taxonomic groups or through time. Finally, disparity measures were calculated and plotted to compare disparity between the different groups, including the sums of variances (differences between morphospace quantity occupied) and sums of ranges (differences in range of occupied morphospace, more sensitive to outliers) (Ciampaglio et al., 2001; Guillerme et al., 2020b). Multiple disparity metrics were calculated as methodological studies have found this to be key to extracting all pertinent information (Ciampaglio et al., 2001; Korn et al., 2013). Pairwise t-tests (alpha level of 0.05 with Bonferroni correction = 0.05/n pairings) were then performed to test for differences in the measured disparities of the groups.

All analyses were run and plots created in RStudio version 2023.09.1+494 (R Core Team, 2023; RStudioTeam, 2020), using the following packages: momocs for geometric morphometric analyses and related statistics (Bonhomme et al., 2014); ggplot2 for plotting (Wickham, 2016); dispRity to calculate disparity measures (Guillerme, 2018). All data, including metadata (specimen accession information, taxonomy, geological age) and cephalon outline landmark coordinates for PCs 1 and 2, are available in Appendix I and online at https://osf.io/vz9a5/. All R code, including a script for data preparation and for all analyses performed, are available online at https://osf.io/vz9a5/.

## Results

### Taxonomic variation in cephalon morphometry

The PCA morphospace with taxonomic order groupings (Fig. 3) shows that, for the most part, trilobite specimens occupy a constrained region of morphospace, with some general differences between order groups and several interesting deviations. Most orders overlap extensively at the centre of the morphospace (Fig. 3A), demonstrating that most trilobite cephala, no matter their taxonomic assignment, display similar morphologies at least in terms of their common outline shapes. However, the orders show differing morphospace occupation in terms of their extremes, and several groups, most notably the Harpida, are offset from the centre of morphospace and display limited overlap with the other orders. The Asaphida cluster tightly within the centre of morphospace (Fig. 3B) as ‘typical trilobites’, with most aulacopleurids clustering also within this region at low values of PC1 and 2. Corynexochida cluster to the left of the centre of morphospace occupation at more negative PC1 values, in a similar region to a dense cluster of Phacopida, though the Phacopida also show dense clustering closer to the morphospace centre (Fig. 3B). Proetida show clustering in an opposing area of morphospace to the Phacopida, with the majority of specimens falling between the two major phacopid clusters and the rest found along more negative PC2 values. It is evident that the Harpida are particularly divergent in cephalon morphometry, with the group forming an almost entirely independent cluster in the most negative region of PC2, though it is interesting that the superficially similar Trinucleida are quite broadly scattered across the morphospace (Fig. 3B).

The Lichida, Odontopleurida and Olenida occupy similar areas of morphospace to each other, mostly at positive PC2 values and reasonably centred along PC1, while the Redlichiida cluster at slightly more positive PC1 values (Fig. 3B). The LDA morphospace shows comparable results to the PCA (Fig. 3C), with most order groups greatly overlapping, the Harpida highly divergent from this general clustering, and some differences in the extremes of various order groupings. This is reinforced by the LDA cross-validation table (Fig. 4), which confirms the Harpida cluster is notably different to the other orders, and demonstrates that the extensive overlap of Phacopida with the other orders would likely cause false predictions for new data points for almost all orders (except Harpida and perhaps Redlichiida).

**Figure 4:**
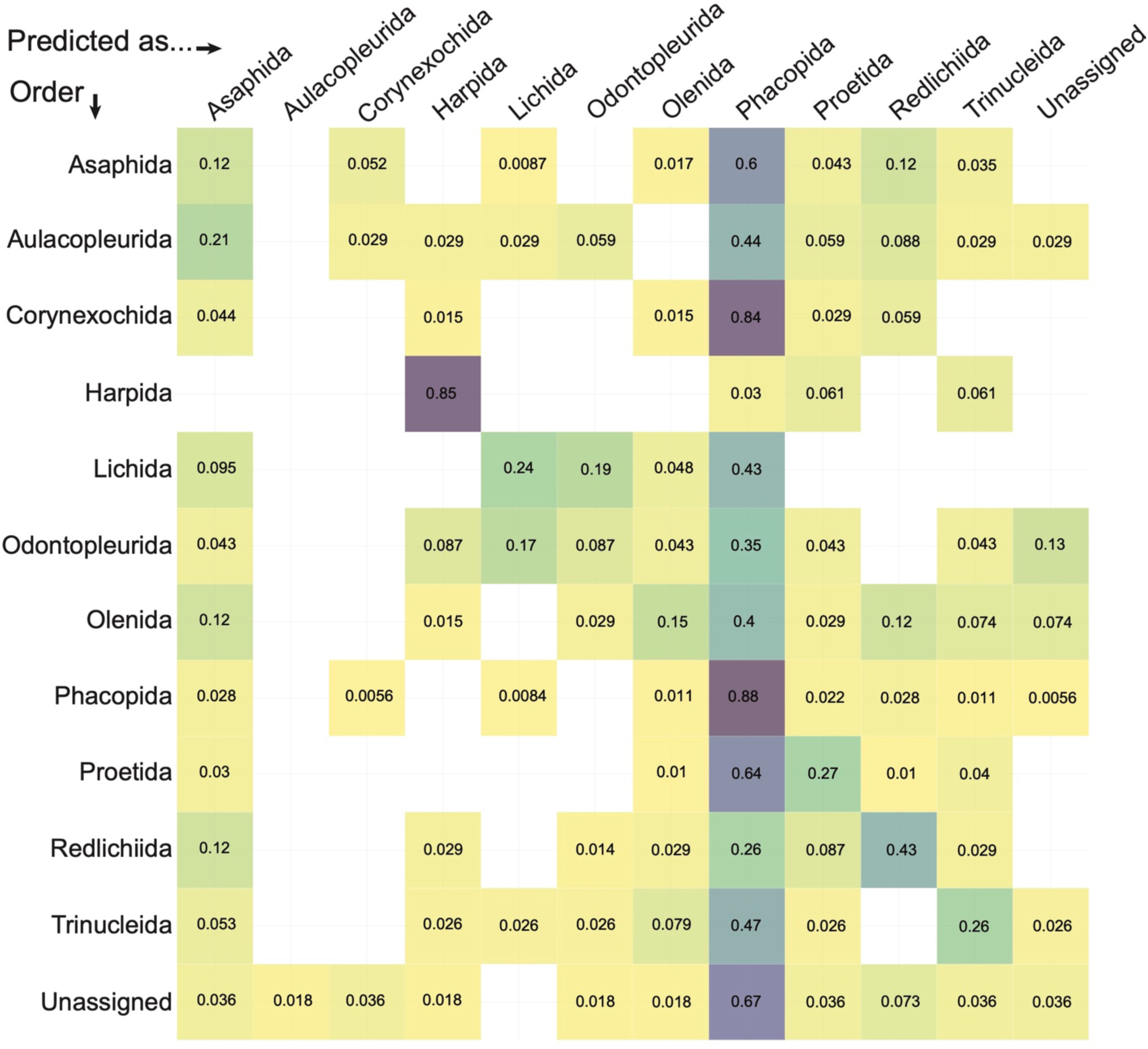
Linear discriminants analysis cross-validation table for data grouped by taxonomic order. Y-axis is the true order of a new data entry, and the x-axis is the order that this new entry would be predicted as being under this dataset.

Pairwise comparisons of the mean cephalon shapes for each order make clear the shape changes responsible for the clustering differences between these groupings (Fig. 5). Harpida shows a highly divergent average cephalon shape, with a much longer (ax.) cephalon with broad and long genal spines. The mean trinucleid cephalon is similar to that of the harpid, but slightly shorter (ax.) and with the genal spines narrower and more distally directed. In contrast, the mean Lichida and Odontopleurida cephala are very shorter (ax.) compared to all other orders. The mean Aulacopleurida, Olenida and Redlichiida cephala are almost identical, as are the Corynexochida and Phacopida cephala (with the Asaphida, the unassigned group and Proetida also alike these; Fig. 5). We can therefore divide the orders into five major trilobite cephalon morphogroups: 1) Harpida; 2) Trinucleida; 3) Lichida–Odontopleurida; 4) Aulacopleurida–Olenida–Redlichiida; 5) Asaphida– Corynexochida–Phacopida–Proetida. These morphogroups are quantitatively supported by the centroid distances between the orders when grouped in PCA morphospace (Fig. 5). The Harpida have high centroid distances to all other groups, and the other morphogroups are reflected in the short pairwise distances between their constituent order centroids. However, the centroid distances to the Trinucleida are not notably high for any order group, and the distance between the Trinucleida and Redlichiida is reasonably low, suggesting similarity between the average cephala of these two groups.

**Figure 5:**
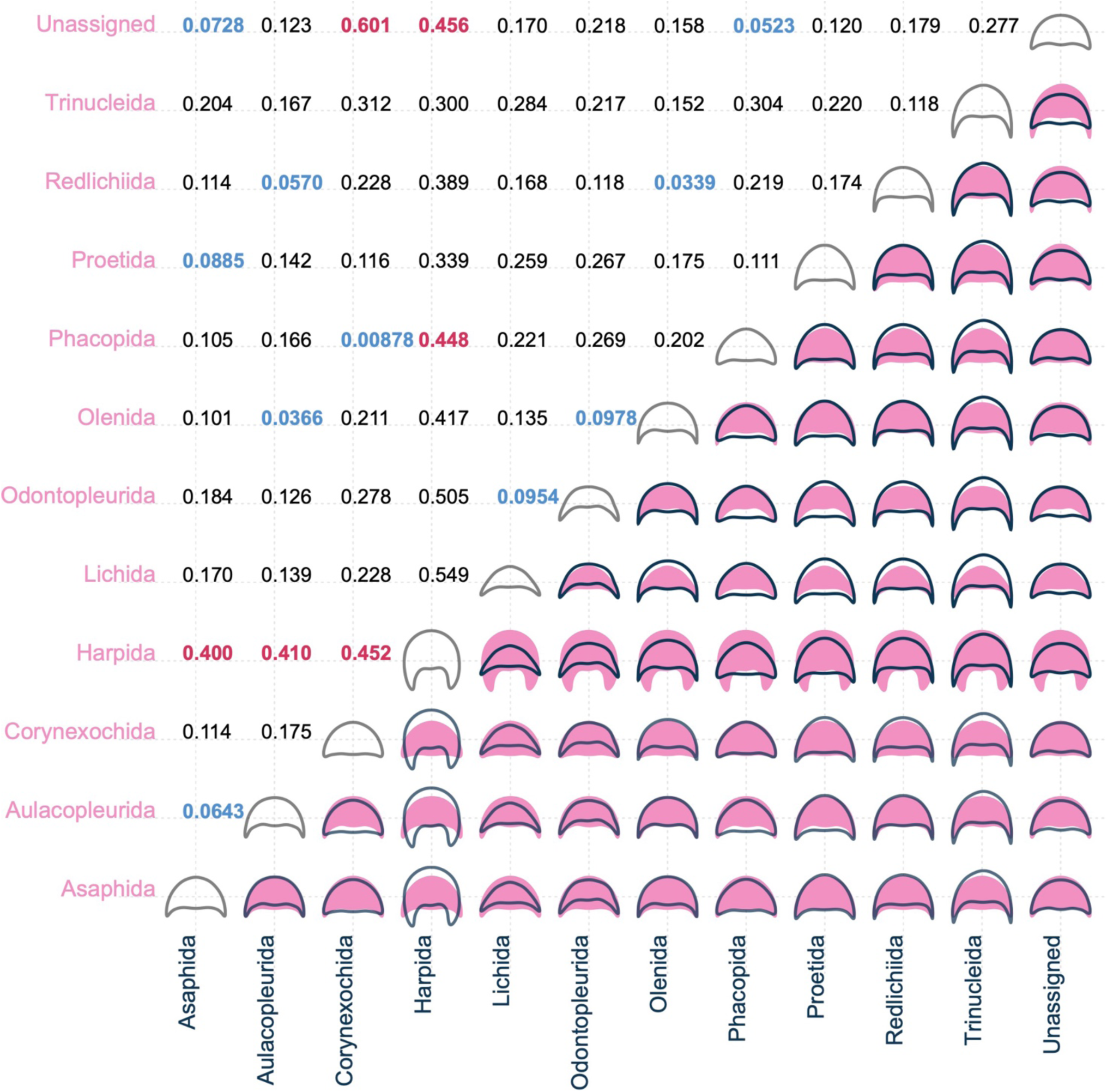
Mean cephalic shape for data grouped within each taxonomic order, compared in a pairwise fashion with the colour of the shape reflecting the colour of its axis (bottom-right side of table). Pairwise centroid distances for taxonomic orders, showing their relative positions in PCA morphospace (top-left side of table); blue figures indicate particularly close pairwise distances (<0.100), and red figures indicate particularly distant pairwise distances (>0.399).

A MANOVA test indicates a significant difference between the distribution of PC scores for the order groupings, suggesting that taxonomy influences cephalic outline data (F = 20.329, p = <2.2✕10^−16^). This is reinforced by many significant pairwise MANOVA tests, testing the null hypothesis that there are no differences in PC score distributions for each pair of orders (see Supplementary I for all test results). Of particular interest are the high F values (all F = >15 and significant p values at p = <7.576✕10^−4^) for the following pairings, which suggest extensive differences in morphospace distributions: the Harpida with all orders; Lichida–Proetida pairing; Odontopleurida–Phacopida; Odontopleurida–Proetida; Olenida–Phacopida; and Phacopida– Redlichiida. These results further support the extensive difference of the Harpida from the other orders, and the existence of the general order morphogroups noted above, particularly the Lichida– Odontopleurida grouping as distinct from the other morphogroups.

The Olenida, Phacopida, Redlichiida and Trinucleida occupy the greatest areas of PCA morphospace (Table 1), with the other orders being comparably constrained (Fig. 3). The Asaphida, Corynexochida, Harpida, Proetida and unassigned group occupy about a half to two-thirds of the morphospace area of the aforementioned groups. The Aulacopleurida, Lichida, and Odontopleurida occupy much less than half of the morphospace area of the groups with the largest areas. Disparity measures generally corroborate this, with Trinucleida, Olenida, Asaphida and Redlichiida showing high sums of variances (Table 1), and the sums of variances comparably low for all other orders (Fig. 6A). The sums of ranges show similar patterns (Fig. 6B), with Trinucleida, Olenida, Redlichiida and Asaphida having high values as for the other measures. However, other orders also show high sums of ranges, most notably Phacopida. The sums of ranges are low for Aulacopleurida, Lichida, Odontopleurida and Proetida, though these sums of ranges are more liable to be impacted by outliers than the sums of variances. Together, these disparity measures suggest high within-group disparity for the Trinucleida, Olenida, Redlichiida and Asaphida, and comparably low within-group disparity for the Lichida and Odontopleurida in particular. The Phacopida interestingly show a comparably large convex hull area and corresponding high sum of ranges, but a relatively low sum of variances, suggesting lower disparity within-group than indicated by their morphospace area occupation. The Asaphida show high disparity metrics for their medium convex hull area, suggesting reasonably high within-group disparity. Pairwise t-tests support these findings (see Appendix III for all test results), suggesting most order groups vary significantly in both disparity measures (Fig. 6). For example, all pairings with the Trinucleida are strongly significant, supporting a statistically higher disparity in this group.

**Figure 6:**
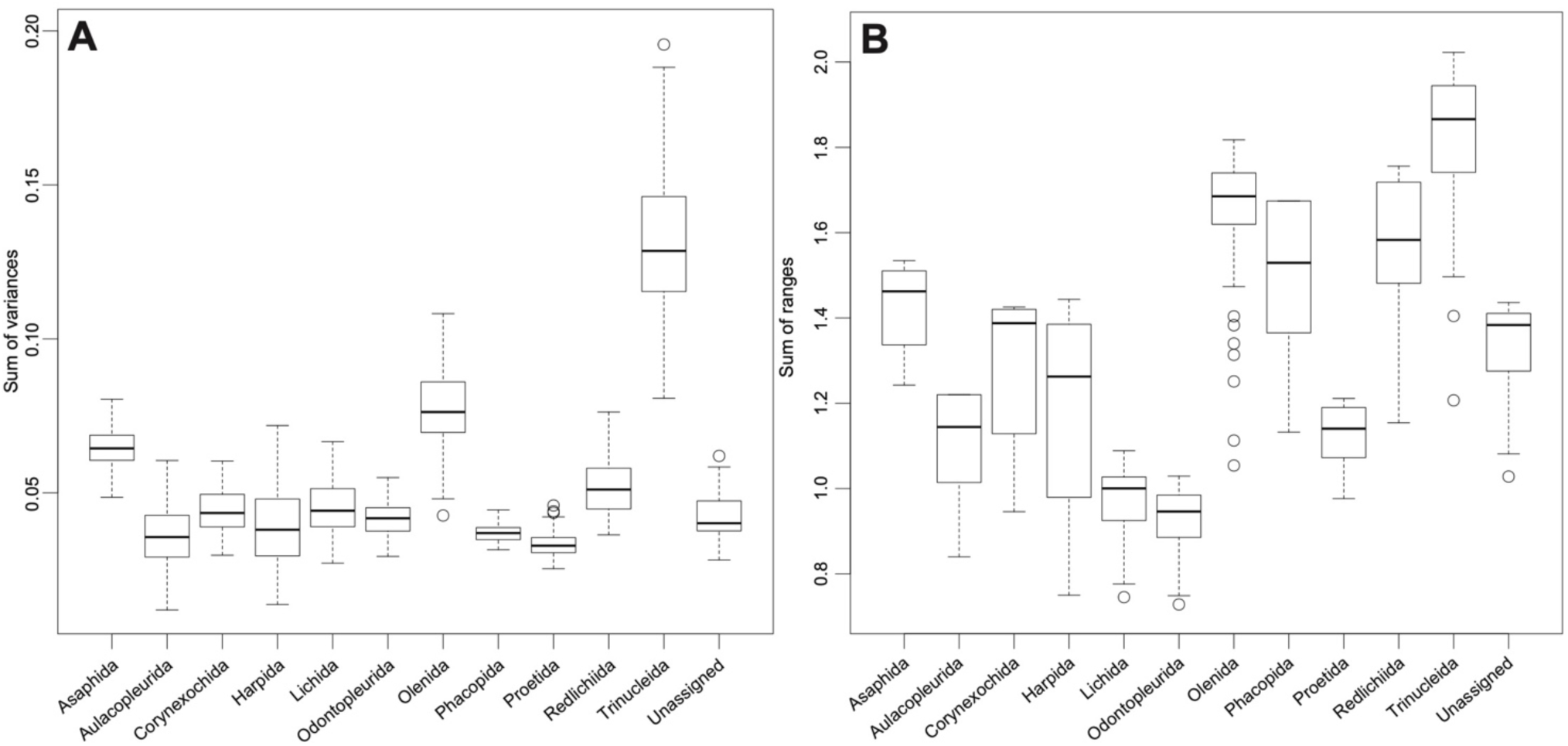
Box and whisker plots showing disparity (space size) within each taxonomic order grouping: sum of variances (A) and sum of ranges (B). Plotted points represent potential outliers.

### Cephalon morphometric variation through geological time

When grouped by geological Period, convex hulls of trilobite cephalic outlines also show considerable overlap in the PCA morphospace (Fig. 7A), though clear differences in the position and relative areas of morphospace trilobites occupied during each Period are evident. Trilobites occupied larger areas of morphospace during the Cambrian to Devonian, with a significant constriction in the Carboniferous and Permian (Fig. 7A and B; Table 2). This constriction occurs towards the centre of the morphospace (and slightly negative on PC2). The PCA morphospace occupancy during the Cambrian is quite diffuse, and only during the Cambrian and Ordovician are the positive extremes of PC1 occupied (Fig. 7B). The Silurian contains some trilobites with negative PC2 values close to the extremes. In comparison, the Devonian shows greater morphospace occupancy to the extremes of PC2, with the PC1 extremes already lost by this time. Interestingly, PCA morphospace occupation during the Devonian appears to comprise three mostly distinct clusters (Fig. 7B); one each to the left and right of the centre of morphospace and extended along PC2, and one cluster at the negative extreme of PC2.

**Figure 7:**
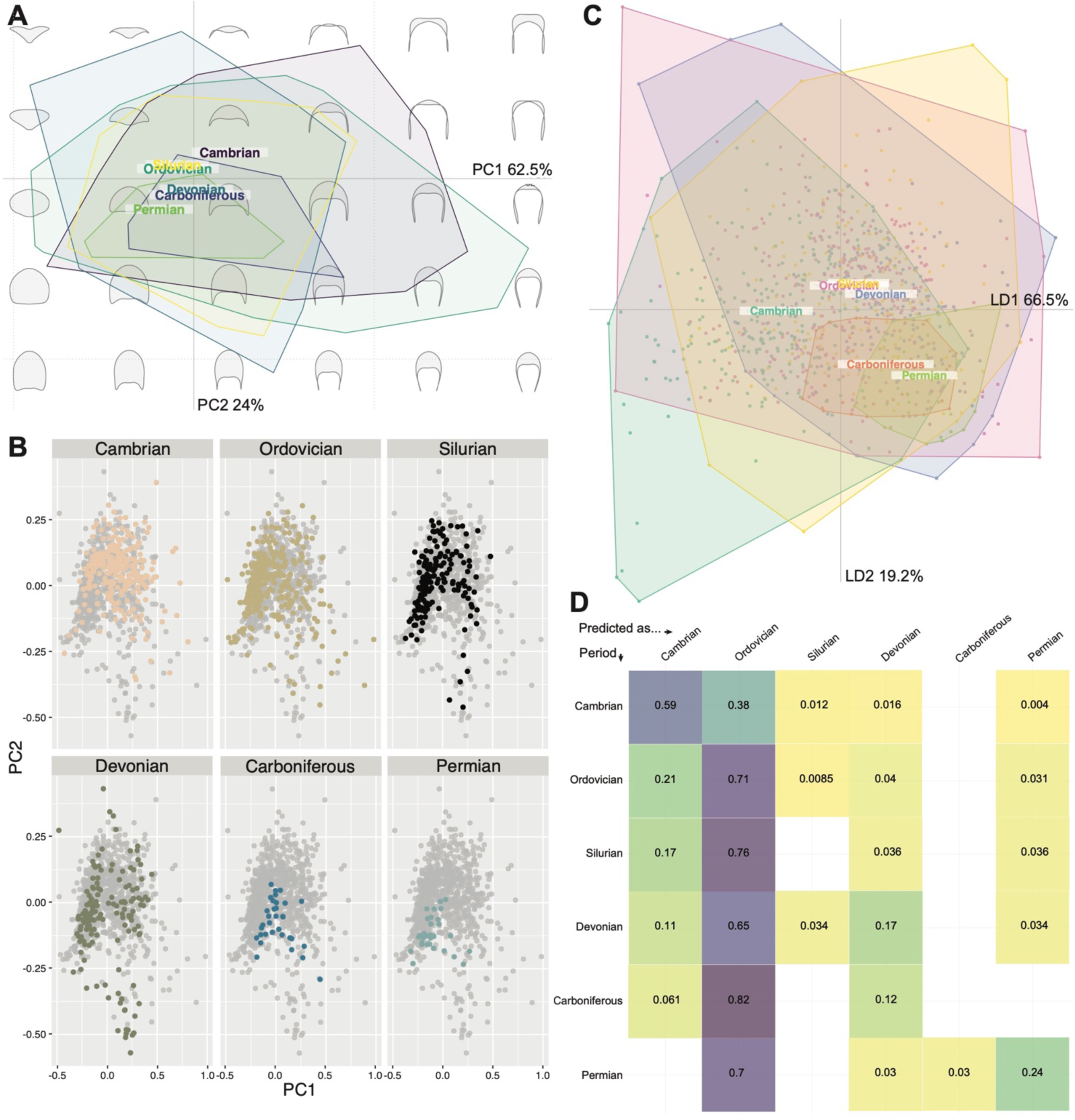
Principal components analysis (PCA) total plot (A), facet wrap (B), linear discriminants analysis (LDA) plot (C), and LDA cross-validation table (D) of trilobite cephalic outline morphometry, grouped by geological Period. A and B visualise the same data, but B has the overlapping convex hull polygons separated out onto individual morphospaces. For D, the y-axis is the true occupied Period of a new data entry, and the x-axis is the Period that this new entry would be predicted as being found in under this dataset. The scree plot for the PCA is the same as that presented in Fig. 3D.

The LDA morphospace (Fig. 7C) differentiates the Cambrian and Ordovician at their extremes, while also reinforcing the centroid of the Cambrian as offset to the other Periods. The centroids of the Carboniferous and Permian are close together but separated from the remaining Palaeozoic Periods. The LDA cross-validation table (Fig. 7D) demonstrates the impact of the extensive overlap of all Periods in morphospace with the particularly broad Ordovician convex hull; data from all Periods except the Cambrian are highly likely to be placed within the Ordovician if a new corresponding data point were added. New data points from the Cambrian are more likely to be predicted correctly than placed in a different Period.

The mean cephalon shapes appear reasonably different for each of the geological Periods (Fig. 8). The Permian mean cephalon is notably long (ax.) with short genal spines, while the Cambrian mean cephalon is much shorter (ax.) with longer and narrower genal spines than for the other Periods.

**Figure 8:**
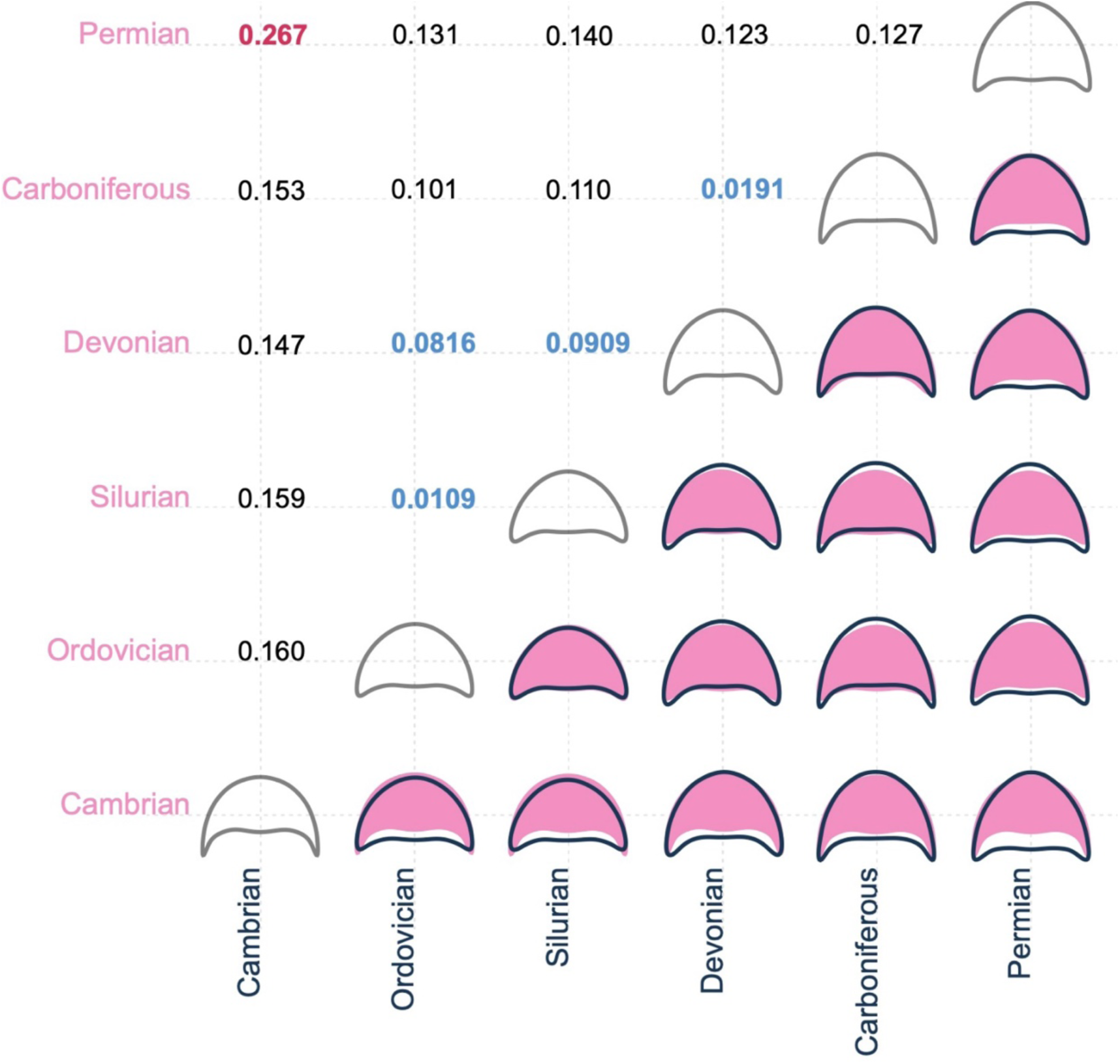
Mean cephalic shape for data grouped within each geological Period, compared in a pairwise fashion with the colour of the shape reflecting the colour of its axis (bottom-right side of table). Pairwise centroid distances for geological Periods, showing their relative positions in PCA morphospace (top-left side of table); blue figures indicate particularly close pairwise distances (<0.100), and red figures indicate particularly distant pairwise distances (>0.200).

Only the mean cephala for the Ordovician and Silurian are similar, and to a lesser extent the Devonian and Carboniferous, though morphogroups are harder to identify than for the order groupings. The pairwise centroid distances are generally low, particularly when compared to those for the order groupings, with the Ordovician–Silurian–Devonian seemingly forming a loose morphogroup with very proximal centroids, and the Carboniferous and Devonian also having centroids close together. The centroid distance between the Cambrian and Permian is notably high (Fig. 8).

Geological age impacted cephalon outline shape, as demonstrated by the significantly different distributions of PC scores for the geological Periods identified by a MANOVA test (F = 9.966, p = <2.2✕10^−16^). Almost every pairing of geological Periods is significantly different to each other following pairwise t-tests, except the Ordovician–Silurian and Carboniferous–Permian (see full test results in Appendix III).

Disparity within the Carboniferous and Permian is low (Table 2; Fig. 9). The Silurian appears moderately disparate based on its relatively high sums of variances and ranges, while disparity is high in all other Periods (Fig. 9). The Devonian is particularly disparate based on sums of variances, and Ordovician based on sums of ranges. All pairwise t-tests were significant, demonstrating that the geological Periods differ in their relative disparities (see Appendix III for all test results).

**Figure 9:**
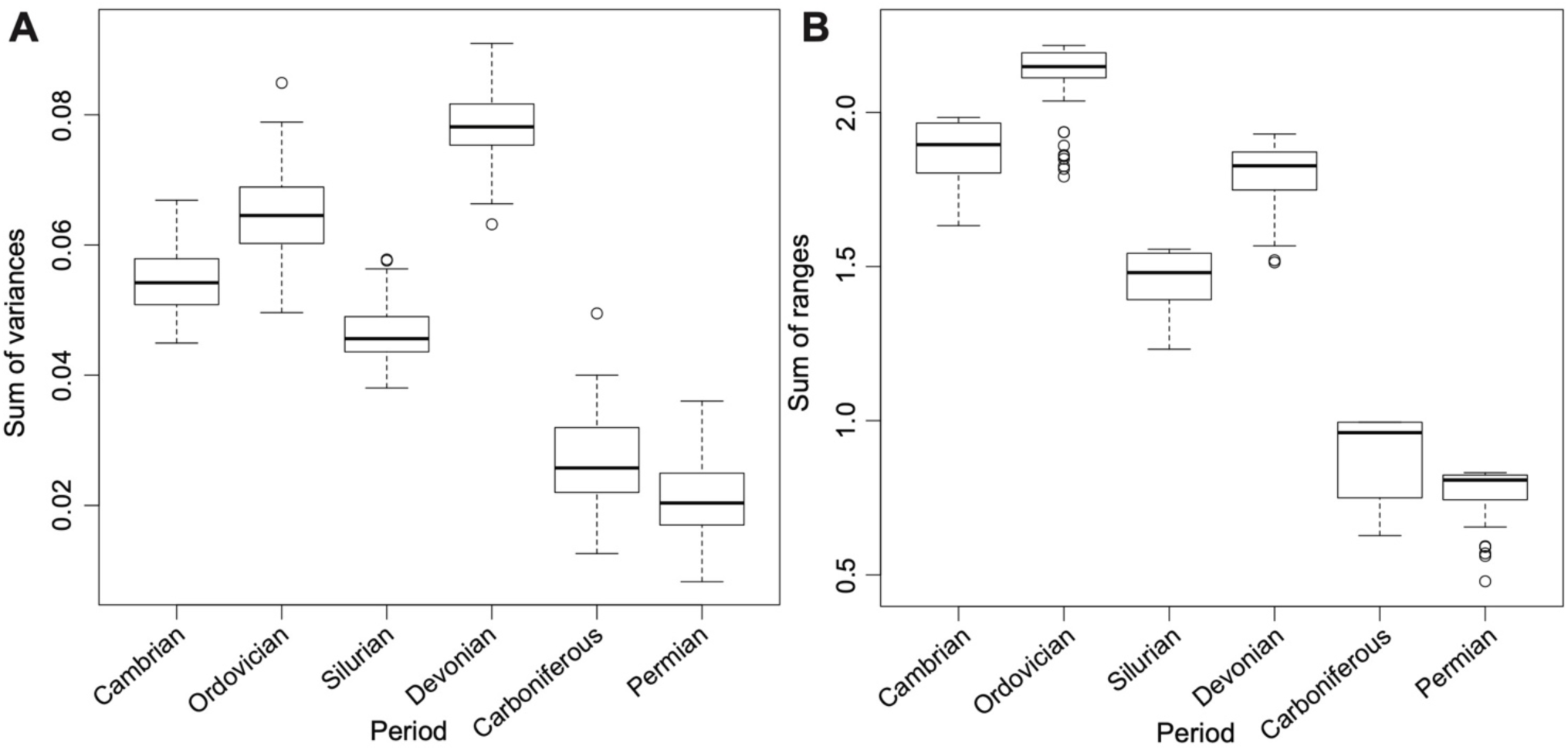
Box and whisker plots showing disparity (space size) within each geological Period grouping: sum of variances (A) and sum of ranges (B). Plotted points represent potential outliers.

### Changing orders through time

Presence and area of morphospace occupation clearly differ for the orders through the Palaeozoic (Fig. 10). There is a radiation indicated by the number of distinct orders between the Cambrian (7 orders) and Ordovician (11 orders), then a decrease to the Silurian and Devonian (7 orders, for both Periods), followed by a severe extinction into the Carboniferous (2 orders) and Permian (1 order remaining). The large area of morphospace occupied in the Ordovician appears partially a result of the evolution of Trinucleida (Fig. 10B). The Harpida occupy a broad range of values along PC2 in the Ordovician (Fig. 10B), which is reduced to a very extreme position at positive PC2 values in the Silurian and Devonian (Fig. 10C and D). Only the Proetida and Aulacopleurida survive to the Carboniferous (Fig. 10E), with the Aulacopleurida also lost by the Permian (Fig. 10F). From their origination in the Ordovician to their demise at the end-Permian, the Proetida are positioned close to the centre of trilobite morphospace occupation.

**Figure 10:**
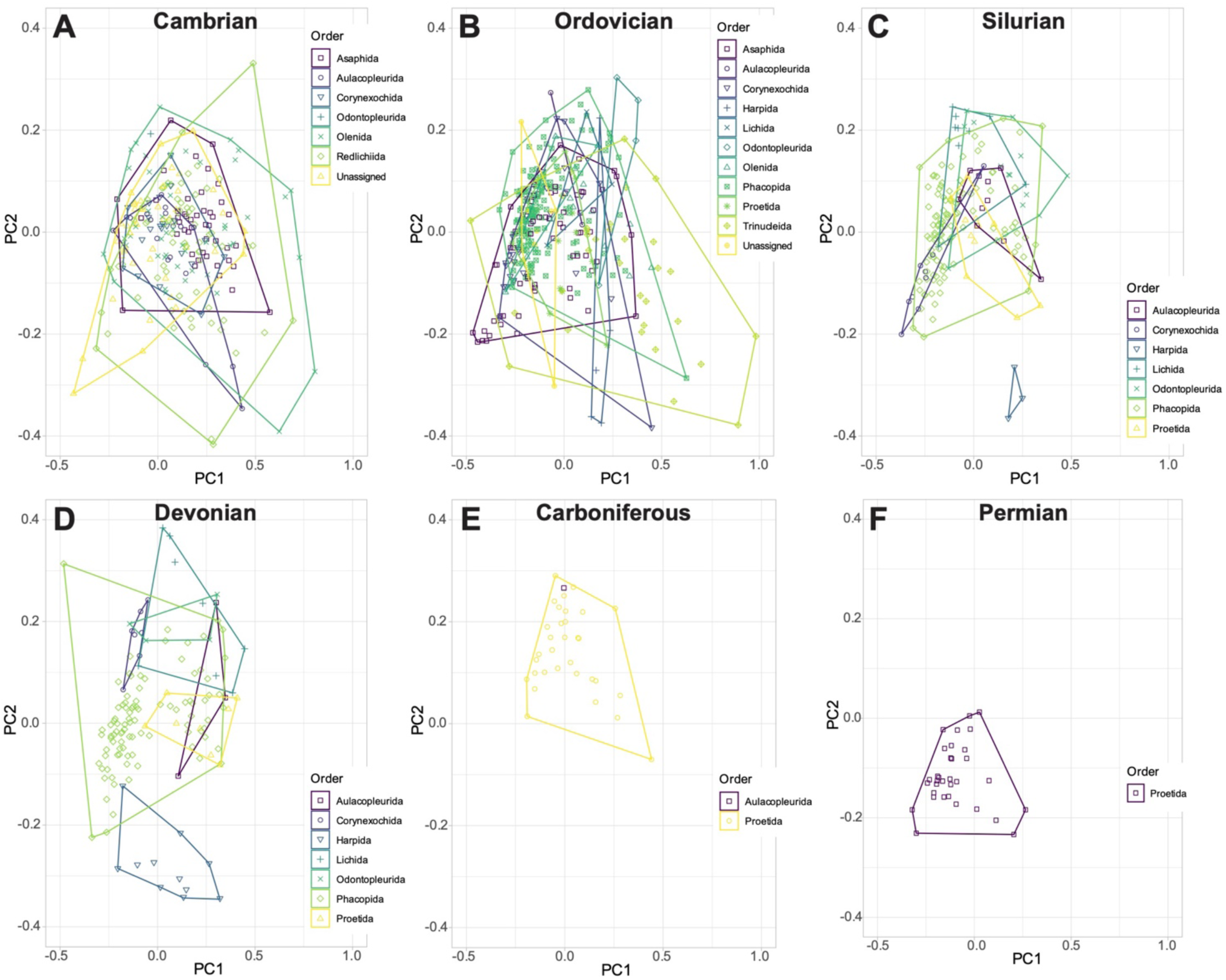
Separate principal components analysis morphospace plots of trilobite cephalic outlines for only specimens present in each Palaeozoic Period. The convex hulls represent the taxonomic orders present in each Period.

### Natural clustering in the dataset

Kmeans clustering analyses produced two major likely hypothetical clustering topologies, which both subdivide the occupied morphospace into seven clusters (Fig. 11). The modelled clusters at the centre, top, and bottom right of morphospace have similar average cephalic morphometries. However, the two topologies differently split the occupation at the left side of the morphospace; one topology splits this side into three clusters (Fig. 11B), and the other into two large clusters (Fig. 11C), with the latter instead splitting the right side of the morphospace into three small, densely packed clusters rather than two.

**Figure 11:**
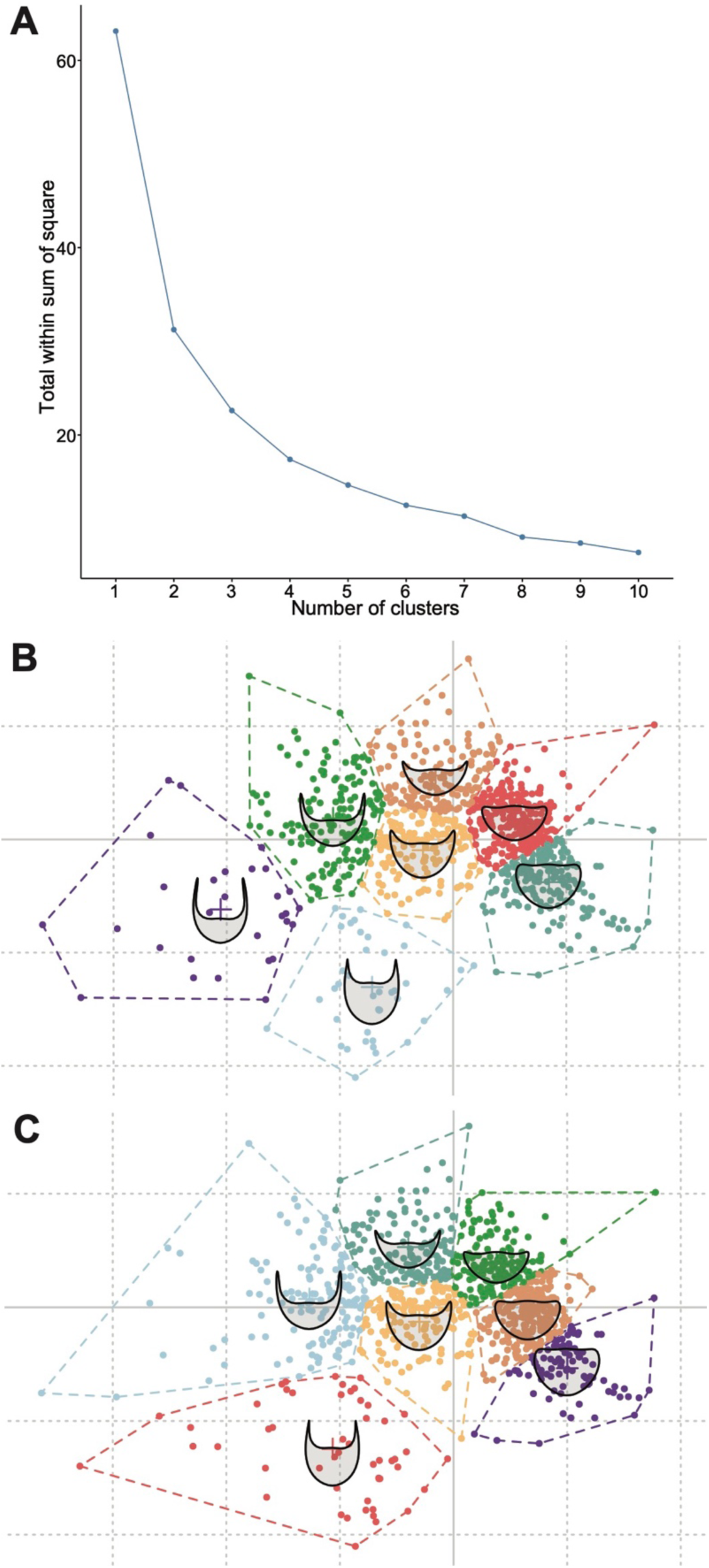
Kmeans analysis displaying the natural clustering inherent in the trilobite cephalic outlines dataset. Elbow plot (A) showing the most suitable number of clusters present in the dataset (7; the point at which there is a plateau followed by a clear down-trending slope on the y-axis). Two, slightly differing, clustering hypotheses using a cluster number of 7 are present (B and C), which closely reflect all possible clustering hypotheses.

Kmeans clustering analyses carried out during each geological Period demonstrate that cephalon morphometry naturally clusters differently through the Palaeozoic (Fig. 12). There is a shift in the dataset from a higher optimal number of clusters (seven clusters; Fig. 11A–C) to a lower number (five clusters; Fig. 11D–F) from the Devonian onwards. These analyses clearly show an overall cephalic shape trend across the Palaeozoic from a higher disparity of forms earlier, with many axially short cephala with long genal spines, to cephalic forms that are more axially long and few with long genal spines. This latter is particularly the case in the Carboniferous and Permian (Fig. 11E and F), where the dataset has only clusters of forms below the PC1 horizontal axis in morphospace. The clusters present are similar between the Cambrian and Ordovician Kmeans analyses (Fig. 11A and B), with slightly more subdivision of the axially long cephala with short genal spines than during the post-Ordovician. The clusters on the left of the morphospace (negative PC1) move more centrally (i.e., away from the extreme left) from the Cambrian to the Silurian, while the clusters in the top and right (positive PC1 and 2) remain approximately consistent into the Silurian. However, by the Silurian the central-most cluster has entirely disappeared, with a new cluster appearing at the bottom extreme of the occupied morphospace (negative PC1; Fig. 11C); this new cluster persists into the Devonian but disappears after this point (Fig. 11D). During the Devonian, several clusters are again positioned centrally on the PC1 axis, with the Carboniferous and Permian both also having a cluster at the very centre of the morphospace (Fig. 11E and F), though the other clusters appear to again move further towards the negative PC2 area of morphospace.

**Figure 12:**
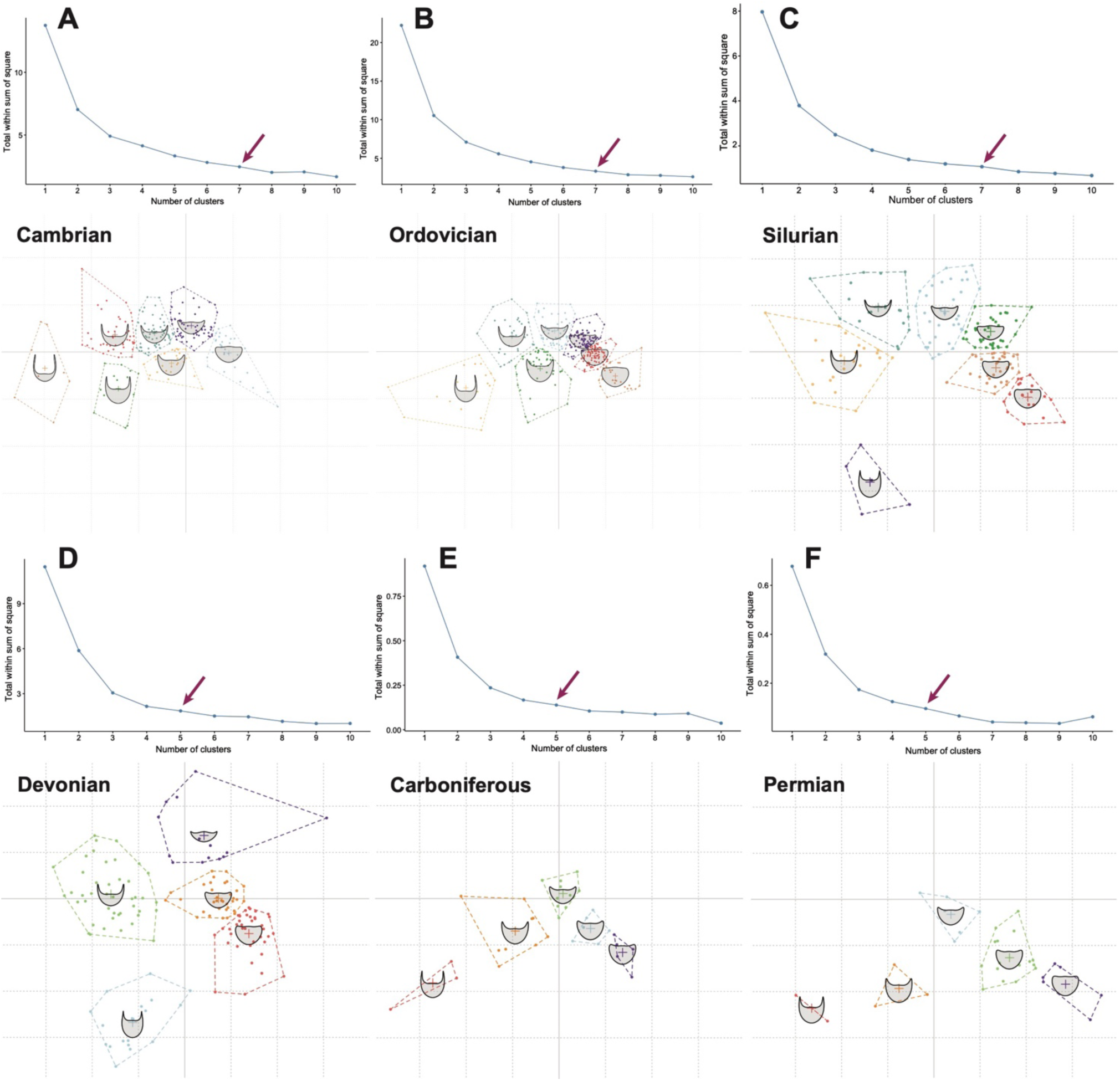
Kmeans analysis displaying the natural clustering inherent in the trilobite cephalic outlines dataset, separated for each geological Period. Both the elbow plot (displaying the most suitable number of clusters) and the clustering hypothesis are presented for each Period. The red arrows display the number of clusters at which there is an observable change (downwards trend) in y-axis steepness.

## Discussion

### Taxonomic signal in cephalon outline morphometry

Cephalic outline shape, alongside the presence and morphology of other cephalic features such as facial sutures, eye ridges, and glabellar furrows, are key for taxonomic descriptions and assignments within Trilobita (Beecher, 1893a, 1893b; Emmrich, 1839; Holmes, 2023; Park and Kihm, 2017; Rasetti, 1948; Salter, 1864; Stubblefield, 1936; Whittington et al., 1997). Indeed, many original species descriptions are based solely on cephala. Although cephalic outline shape reflects only part of the complexity of trilobite morphology, and there is broad overlap of many orders in the central part of the morphospace (Fig. 3), a clear taxonomic signal is present in our morphospace. This can most clearly be seen by the separation of Harpida from the other orders (Figs 3–5), a result of their unique morphology comprising a broad brim, high axial length, and elongate flattened genal projections. Trinucleida were also found to differ significantly from other orders in the dataset, occupying a broad area of morphospace distinct from harpids (Fig. 3), likely as a consequence of their thinner genal spines and differences in the outline shape of the fringe.

The relative positions of orders in our morphospace provide support for some hypotheses for high-level evolutionary relationships and historical taxonomic assignments that have been notably difficult to draw lines between. In particular, a close phylogenetic relationship between the Lichida and Odontopleurida (Fortey, 2001; Thomas and Holloway, 1997), though other works disagree (Adrain, 2013), and the almost identical average cephalic outline of Redlichiida and Olenida, suggesting a close relationship between the two near the origin of trilobites in morphospace (g.g., Bault et al., 2022). Though cephalic outline is clearly unable to elucidate problematic ordinal-level relationships, such as for Trinucleida, which is suggested to have a closest similarity to the unassigned, previously ptychopariid, group (Bignon et al., 2020).

Distinct differences between some major orders, in addition to the Harpida and Trinucleida, are also clear from the analyses. Phacopida and Proetida cluster in somewhat opposite patterns in morphospace, occupying the same area on PC1 but with Proetida more offset towards negative PC2 values (an axially longer cephalon; Fig. 3). This may reflect the differences in the orders’ broad-scale evolutionary trajectories, with phacopids being more diverse earlier in the Palaeozoic and proetids later, resulting from apparent turnover events (e.g., Bault, 2023). Proetida is notably constrained at the centre of the morphospace (Fig. 3), suggesting little late-Palaeozoic exploration of extreme cephalic morphologies and potentially a constriction in viable forms through time (Fig. 10). This presumably reflects external selection pressures on proetids during the Carboniferous and Permian as the only trilobite order surviving into the Permian, over their decline to eventual extinction (e.g., Adrain, 2013; Bault, 2023; Bault et al., 2022; Fortey, 2001). Lerosey-Aubril and Feist (2012) found that proetids in the late Palaeozoic experienced many separate extinction and radiation events, though our dataset suggests this is not reflected in their cephalic shape disparity averaged across this time, and our time bins are too coarse to provide the required resolution to detect both extinctions and radiations within geological Periods. Phacopida also comprises two major clusters either side of the centre of morphospace, with the group’s major differentiation being along the PC2 axis (primarily axial length of cephalon, with other cephalic margin features being reasonably consistent). Results such as this bimodal clustering suggest interesting underlying differences in morphospace occupation at lower taxonomic levels.

Redlichiida and Olenida occupy a reasonably wide area of morphospace despite the former living only during the Cambrian and the latter the Cambrian and Ordovician (Monti and Confalonieri, 2019), a more narrow geological range than for many other orders. The Olenida and Redlichiida may have both originated effectively contemporaneously in the early Cambrian, potentially reflecting a close evolutionary relationship, and this could explain their comparable morphospace occupation (Fig. 3) and almost identical average cephalic outline shape (Fig. 5). For the Redlichiida, the broad morphospace occupation might indicate the early, labile exploration of trilobites in cephalon shape during their evolution, particularly for the early Cambrian Olenellina (Lieberman, 1998), and when they might have had fewer competitors for ecological niche space that was continually partitioned (Conway Morris and Fritz, 1984; Na and Kiessling, 2015), prior to constriction to a more fixed form. However, the redlichiid cephalon is generally centred within the morphospace, potentially suggesting less exploration of overall form than in orders living during the Ordovician to Devonian (Figs 3 and 7). Similarly, Asaphida occupy a relatively broad area of morphospace in both the Cambrian and Ordovician, but, as suggested by previous studies, suffered greatly in the end-Ordovician extinctions (Fortey, 2001; Speyer and Chatterton, 1989), hence their post-Ordovician disappearance in the dataset.

Some orders with lower sample sizes occupy smaller areas of morphospace, such as the Aulacopleurida, Lichida, and Odontopleurida (all <40 specimens; Table 1). However, there is not a consistent link between sampling and morphospace area occupied; several orders with notably large sample sizes occupy smaller areas of morphospace, for example, the Asaphida and Proetida (Table 1, Fig. 3). This likely reflects the limited cephalon outline disparity of these orders (Fig. 6), for their diversities, compared to other orders. The disparity results also do not correlate well with sampling, for example, disparity measures are notably high for the Trinucleida, despite its low sample size (Table 1, Fig. 6). Additionally, the taxonomic trends presented here are not heavily biased by sampling, supported by comparison of bootstrapped disparity metrics, and thus likely represent real signals in the data.

However, despite the noted differences in morphospace occupation, the extensive overlapping of almost all orders (Fig. 3) indicates high similarity of cephalon outline shape within trilobites, if these taxonomic assignments are taken as reflecting true phylogenetic relationships. As evidenced by the LDA results (Fig. 4), cephalon outline shape alone lacks predictive power at the order level, as only harpids can be correctly predicted with any likelihood. This is further supported by the Kmeans clusters, which directly interrogated the stochastic clustering of the data, without the overlain presupposed clustering enforced by taxonomy. The Kmeans cluster in the lower part of the space (particularly in Fig. 11B) largely reflects the location of the Harpida in morphospace (Fig. 3B), and these clusters thereby show similar average morphologies. However, despite some minor similarities, for example, the Asaphida has a similar average morphometry to the central Kmeans cluster, most other orders do not obviously align with any of the Kmeans clusters.

### Two peaks in morphospace occupation in the Palaeozoic

Trilobite cephalic outline shape varied across the Palaeozoic, both in terms of area of occupied morphospace and average position within morphospace (Figs 6, 7). Overall, there is a high initial area of morphospace occupation in the Cambrian, which then increases to its maximum in the Ordovician, with a second, slightly lower, peak in the Devonian. Following the Devonian, morphospace occupation is much lower in the Carboniferous and Permian. An Ordovician peak in morphospace occupation differs from the previous study using cephalic outlines, where a peak in occupation was found in the Cambrian (Suárez and Esteve, 2021), though is consistent with older studies using cranidial outlines (Foote, 1993b, 1991). The second, smaller, peak in morphospace occupation during the Devonian was recovered by both sets of studies (Foote, 1993b; Suárez and Esteve, 2021). The post-Devonian morphospace constriction is reflected in the Kmeans clustering present in the dataset across the Devonian–Permian (Fig. 12); the optimal number of Kmeans clusters decreases following the Silurian due to the restriction of specimens to the central-lower part of the morphospace, with fewer clusters reflecting the loss of more extreme morphologies. Overall, the results suggest a picture of changing cephalon morphometry through the Palaeozoic from axially shorter with longer genal spines, to axially longer with shorter genal spines.

Interrogation of the relative position of centroids in the Ordovician and Devonian (when occupied morphospace peaks) compared to the previous Periods (Cambrian and Silurian respectively) suggests distinct underlying mechanisms for the expansion in morphospace area for each Period (Fig. 7A, B). During the Cambrian, cephalic forms mostly occupy the central region of morphospace, then from the Ordovician trilobites split away from the centre along PC1 with major clusters just to the left and right of the centre (supported by the differentiation of the Cambrian and Ordovician by the LDA; Fig. 7C). These discrete clusters and differences in centroid position suggest that different trilobite forms existed in the Cambrian compared to the post-Cambrian. This lateral shift in morphospace occupation reflects the well-established cryptogenesis problem within trilobites, where morphological similarities that link Cambrian to post-Cambrian trilobites have been difficult to identify (recently reviewed by Paterson, 2019, see also Fortey, 2001). Notably, the redlichiids lived only during the Cambrian, with a suborder of redlichiids, the Olenellina, having particularly different cephalic morphologies to post-Cambrian trilobites, such as a lack of facial sutures but presence of a ventral moulting suture (Esteve et al., 2021; Henningsmoen, 1975; Hughes, 2007). Importantly, the Ordovician saw the origins and radiations of a range of new trilobite orders, families and genera (Adrain et al., 1998; Paterson, 2019). All major trilobite orders were thereby established by the late Ordovician (Fig. 10B), establishing the body plans that were embellished upon for the remainder of the Palaeozoic.

The Ordovician saw trilobites expand not just in terms of morphological differences (morphospace space size) but also in terms of what the ‘average trilobite’ looked like (morphospace centroid position). Indeed, this shift in centroid position from the Cambrian to the Ordovician is the largest of all trilobite centroid shifts in the Palaeozoic (Fig. 7), indicating that trilobites were radiating into a new set of niches at this time (e.g., Droser & Finegan, 2003; Serra et al., 2023). A contemporaneous taxonomic and ecological radiation is not surprising, as the expansion of trilobites into new niches will have been facilitated by new morphological innovations, which in turn allow taxonomic groups to be diagnosed. This broadly coincides with the Great Ordovician Biodiversification Event (GOBE), later stages of the plankton revolution (Servais et al., 2016, 2010), and a time when trilobites explored new ecologies as juveniles and adults, with corresponding impacts on cephalic morphology. Ecological diversity in trilobites has been suggested to peak during the Ordovician (Fortey & Owens 1990). For example, several groups independently evolved planktic larvae towards the late Cambrian and early Ordovician (Chatterton & Speyer 1989; Laibl et al., 2023), trilobites are thought to have occupied a range of levels in the water column during the Ordovician (Esteve and López-Pachón, 2023; Fortey, 2014, 1985; McCormick and Fortey, 1998), and some species expanded into deeper and tropical environments (Hopkins, 2014). The shape of the cephalon would have played an important role for moving and feeding within the water column, under conditions very different to those encountered by benthic forms (e.g., Fortey 1985; Esteve and López-Pachón, 2023). Closer to the seafloor, ecological pressures such as predation have been suggested to have led to embellishments in the trilobite exoskeleton, including the cephalon (Hughes 2007).

In contrast, there is no evident shift in centroid position from the Silurian to the Devonian, despite a relatively larger increase in the area of morphospace occupied during this transition (compared to the Cambrian–Ordovician; Fig. 7). This expansion in morphospace is likely driven by the relatively higher diversity of trilobites within the same orders in the Devonian compared to the Silurian (in our dataset; Bault et al., 2022), rather than specialisation to new niches. Support for this interpretation comes from visual inspection of the morphospace, with expansion in all directions (Fig. 7B), and quantification of the morphospace occupation areas (Table 2). Expansion in morphospace area occupation by trilobites at this time was thereby driven by morphological innovation by existing orders. This can be seen most clearly in the Harpida region of the morphospace (Fig. 10C, D), with the harpids innovating following decline during the Silurian, as observed by Beech and Lamsdell (2021). Lichida and Phacopida also expand their occupation of morphospace in the Devonian, however, other trilobite orders (e.g., Aulacopleurida, Corynexochida, Proetida) did not contribute to this increase in morphospace occupation (Fig. 10C, D), demonstrating heterogeneity in response.

Constrictions in morphospace occupation (Ordovician to Silurian; Devonian to Carboniferous; Carboniferous to Permian) are also not associated with large shifts in centroid position (Fig. 7). This indicates that reduction in morphospace occupation is associated with declining diversity of trilobites (Fig. 10), but no clear extinction specificity can be observed. Previous studies have linked selectivity for extinction to the ecology of trilobite larvae (Speyer and Chatterton, 1989), however, we do not see support for this at the cephalon outline level of sampled adults. Similarly, the global eustatic perturbations, anoxia of bottom waters, and broad environmental changes associated with Devonian extinction events, which likely led to the extinction of deep-water groups (e.g., Lerosey-Aubril & Feist, 2012) suggest some extinction selectivity, but the data here, at this temporal resolution, do not find this. The lack of a selective extinction signal is likely the result of several factors. Firstly, selective extinction of trilobites with specific juvenile ecologies did not necessarily translate to selectivity in adult cephalic shape. Secondly, selective changes may have applied only to specific internal cephalic structures, such as the parallel eye reduction and facial suture shape change visible in many late Devonian trilobites as a response to global sea level changes (e.g., Feist, 1991; Feist & McNamara, 2013). Lastly, and most importantly, the data sampling presented is likely too coarse. Selectivity of extinctions will be hidden by post-extinction radiations of trilobites reoccupying morphospace areas vacated by extinct groups, and without sampling at lower taxonomic levels this will be obscured. This is presumably the case for Carboniferous proetids, where selective family extinction and radiation has been obfuscated. Following the end-Devonian extinction, proetids expanded into areas of morphospace it had not previously occupied (Fig. 10D– F), including a dense band close to an area originally dominated by Phacopida and a space proximal to the positive extreme of PC2 previously occupied by Harpida. This increase in disparity reflects an increase in generic level diversity that is decoupled from the stagnant order level diversity. In turn, this demonstrates the capacity that proetids had to adapt following the Devonian mass extinctions (Lerosey-Aubril and Feist, 2012), potentially facilitated by the removal of harpids and phacopids. This proetid expansion was likely driven by high origination rates in the early Carboniferous, however, this origination rate dropped dramatically in the Viséan (c. 350 Ma) and did not recover (Lerosey-Aubril and Feist, 2012). The continued winnowing of trilobite diversity, and with it disparity, was likely the result of major environmental changes associated with the icehouse Carboniferous and subsequent formation of Pangaea (Lerosey-Aubril and Feist, 2012). Further global environmental changes occurred in the Permian, and their impact on trilobite diversity and disparity and are reflected by the further constriction of morphospace area occupation at the Carboniferous–Permian boundary, and eventual extinction of trilobites.

Measures of morphospace occupation can be influenced by sample size, and this is potentially problematic for quantitative comparisons of morphospace area later in the Palaeozoic, with the two Periods with the lowest morphospace areas occupied and lowest disparities being those with the smallest sample sizes (Fig. 7, Table 2). However, as for orders, morphospace area occupied does not consistently track with sample size, for example, sample size is higher in the Silurian than the Devonian, but morphospace area is higher for the Devonian (Table 2). Further, the sampling does reflect the much lower species richness in the Carboniferous and Permian resulting from the prior extinctions of almost all trilobite orders (e.g., Bault et al., 2022; Fortey, 2001; Lerosey-Aubril & Feist 2012).

Despite these evident trends in cephalic outline disparity and trilobite diversity, the predictive power of cephalon morphometry for geological Period is generally poor (Fig. 7D), as for the taxonomic orders. Only the Cambrian and Ordovician groupings were predictive, with the average Ordovician cephalon being axially longer, and the Cambrian cephalon having longer genal spines (Fig. 8). Thus, this large and broadly sampled dataset suggests that cephalon outlines could not generally be used to predict the order assignment or the occupied geological Period of an unknown trilobite.

## Conclusions

A large dataset of almost 1000 2D trilobite cephalon outlines was used to explore the extent of trilobite occupation of morphospace, and to determine whether this cephalic morphometry was associated with order assignment or geological Period occupied. The data show significant differences in morphospace occupation and disparity, supporting a reasonably strong taxonomic signal and changes in disparity through the Palaeozoic. Two peaks in trilobite cephalic disparity are observed, attributed to the Ordovician and the Devonian. Importantly, the expansion of morphospace occupation in the Ordovician was presumably due to the occupation of new niches by trilobites (linked to the origin of all trilobite orders without Cambrian origins), contrasting it to the second peak of morphospace occupation in the Devonian, which was likely linked solely to within-order taxonomic diversification. However, due to extensive overlapping of morphospace occupation, clustering analyses indicate that the order or Period assignment of a new unknown trilobite would be almost impossible to predict with any accuracy based on 2D cephalic morphometry. Only the cephalic outlines of Harpida, and of Cambrian and Ordovician trilobites, are predictive.

## Author contributions

**Harriet B. Drage:** Conceptualisation, Methodology, Software, Validation, Formal Analysis, Investigation, Data Curation, Writing – Original Draft, Writing – Review & Editing, Visualisation, Project Administration. **Stephen Pates:** Conceptualisation, Methodology, Software, Validation, Formal Analysis, Investigation, Data Curation, Writing – Original Draft, Writing – Review & Editing, Project Administration.

## Supporting information

Appendix III

## Acknowledgements

We are grateful to Jana Bruthansová (National Museum Prague, Prague), Richard Howard and Zoe Hughes (Natural History Museum UK, London), Matt Riley (Sedgwick Museum, Cambridge), Mónica M. Solórzano Kraemer (Senckenberg Museum, Frankfurt), and Julien Kimmig (Staatliches Museum für Naturkunde Karlsruhe) for providing access to specimens at their respective institutions. In addition, we thank all the museums, volunteers and staff who uploaded images of trilobites to iDigBio, which facilitated data collection for this research during periods of remote working during early 2021. HBD was funded under a Swiss National Science Foundation Sinergia grant (198691). SP was supported by a Herchel Smith Postdoctoral Fellowship (University of Cambridge).

